# Leukemia Risk Factor ARID5B Coordinates HDAC-Mediated Transcriptional Repression

**DOI:** 10.1101/2025.10.17.683040

**Authors:** Ana P. Kutschat, Fabian Frommelt, Brianda L. Santini, Sophie Müller, Paul Batty, Gerlinde Karbon, Giulio Superti-Furga, Davide Seruggia

## Abstract

Multiple genetic association studies linked variants at *ARID5B* with predisposition to B-cell derived acute lymphoblastic leukemia (B-ALL) in children. Still, the molecular function of ARID5B remains largely uncharacterized. Here, we employ a combination of proteomics, genomics and transcriptomics to describe the molecular mechanisms of ARID5B. We identify that ARID5B interacts with MIER1, C16ORF87, HDAC1 and HDAC2 forming a chromatin repressor complex. By CUT&RUN, we mapped ARID5B binding in active regions of the genome, tethering HDAC1 and HDAC2 to distal regulatory elements and promoters. Genes actively repressed by the ARID5B repressor complex are involved in B cell proliferation and B cell-specific signaling. Together, we describe how ARID5B assembles into a repressor complex and regulates B cell-specific processes. Understanding its molecular mechanism will help elucidating how non-coding germline variants at ARID5B predispose to B-ALL.

## Introduction

Germline non-coding variants are known risk factors of pediatric B-cell acute lymphoblastic leukemia (B-ALL)^1–13^. Importantly, some risk variants are in eQTL and are associated with lineage-specific transcription factors, such as *IKZF1* and *CEBPE*^1,2^, suggesting that dysregulation of lineage-specific transcriptional programs contributes to the predisposition to B-ALL. Among the strongest associations are risk variants localized in intronic regions of *ARID5B*^1,2,14^ (AT-Rich Interaction Domain 5B), a DNA-binding protein whose function is not clearly understood. Furthermore, individuals carrying the risk allele or a heterozygous deletion of *ARID5B* oftentimes express lower levels of the gene^14,15^, highlighting the importance of ARID5B in the onset of leukemia.

Recently, inactivation of ARID5B in the mouse was described to affect B cell development promoting an accumulation of pre-B cells and decrease in the immature B cell population in the bone marrow^16^. In line with these findings, ARID5B overexpression in mice leads to a decrease of the pre-B cell (Hardy fraction D) population in the bone marrow^17^. ARID5B was further shown to promote tonic BCR signaling, to limit cell proliferation and regulate fatty acid uptake and metabolism in B cells^16,17^.

*ARID5B* encodes for two isoforms (short and long), with both isoforms having an ARID (AT-Rich Interaction Domain), responsible for DNA-binding, and only the long isoform containing a BAH (Bromo Adjacent Homology) domain^18^. Disruption of the DNA binding of ARID5B has been shown to lead to the de-repression of transcriptional regulators of adipocyte thermogenesis^19^. In turn, the BAH domain of ARID5B is predicted to mediate protein-protein or protein-nucleic acid interactions on chromatin^18^. In fact, ARID5B has been suggested to interact with the histone demethylase PHF2, regulating the expression of metabolic genes in the liver^20^ and genes involved in chondrogenesis^21^.

Thus, while experiments in mice^16,17^ highlighted the implication of ARID5B in coordinating key cellular processes during B cell development, especially at the pre-B cell stage, the molecular mechanisms by which it regulates these processes is still unknown. In this study, we used proteomics to map the ARID5B protein network and identified transcriptional repressors HDAC1, HDAC2, MIER1 and C16ORF87 as main interactors. By targeting ARID5B protein domains, we mapped the BAH domain as the main interface mediating the ARID5B-HDAC interaction. Upon inactivation of ARID5B, we observed a significant loss of HDAC1 and HDAC2 on chromatin and concomitant gain in H3K27ac, suggesting that ARID5B is required to recruit HDAC1 and HDAC2 to chromatin. Gain of H3K27ac in these regions upon HDAC inhibition further confirms that these loci are directly repressed by HDAC1 and HDAC2. Finally, transcriptomics revealed that ARID5B and HDAC regulate genes involved in B cell proliferation and B cell-specific signaling.

## Results

### ARID5B assembles in a complex with HDAC1, HDAC2, MIER1 and C16ORF87

To map the protein-protein interactions of ARID5B, we conducted affinity purification followed by mass spectrometry (AP-MS) (**Fig. 1A**). For this, we overexpressed the long isoform of the murine ARID5B tagged with a FLAG-AviTag in HEK293T cells expressing the E. Coli biotin ligase BirA (**Fig. S1A-B**). In the presence of biotin, existing in low amounts in the cell culture medium, BirA biotinylates the AviTag, facilitating the affinity purification of ARID5B and its interactors using streptavidin beads, which can be quantified by mass spectrometry^22^. Notably, PHF2 was not identified, and consequently no interaction of ARID5B with PHF2 was recovered, contrary to previous reports^20,21^ (**Fig. 1B**; **Table S1**). Instead, we detected significant interactions with HDAC1, HDAC2, MIER1 and C16ORF87. While the latter is an uncharacterized protein containing a zinc-ribbon-domain (PFAM PF10571), HDAC1 and HDAC2 are components of six transcriptional repressor complexes^23,24^, one of them also including the ELM-SANT-domain containing protein MIER1^23,25^. Importantly, ARID5B is identified in the reciprocal affinity enrichment of HDAC1, HDAC2, MIER1 and C16ORF87 in publicly available protein-protein interaction (PPI) databases. A protein interaction network based on PPI-databases covers 18 out of all 20 potential edges/ interactions among ARID5B and its interaction partners (**Fig. S1C, Table S2**), suggesting that ARID5B associates with HDAC1, HDAC2, MIER1 and C16ORF87 into a protein complex.

**Figure 1:**
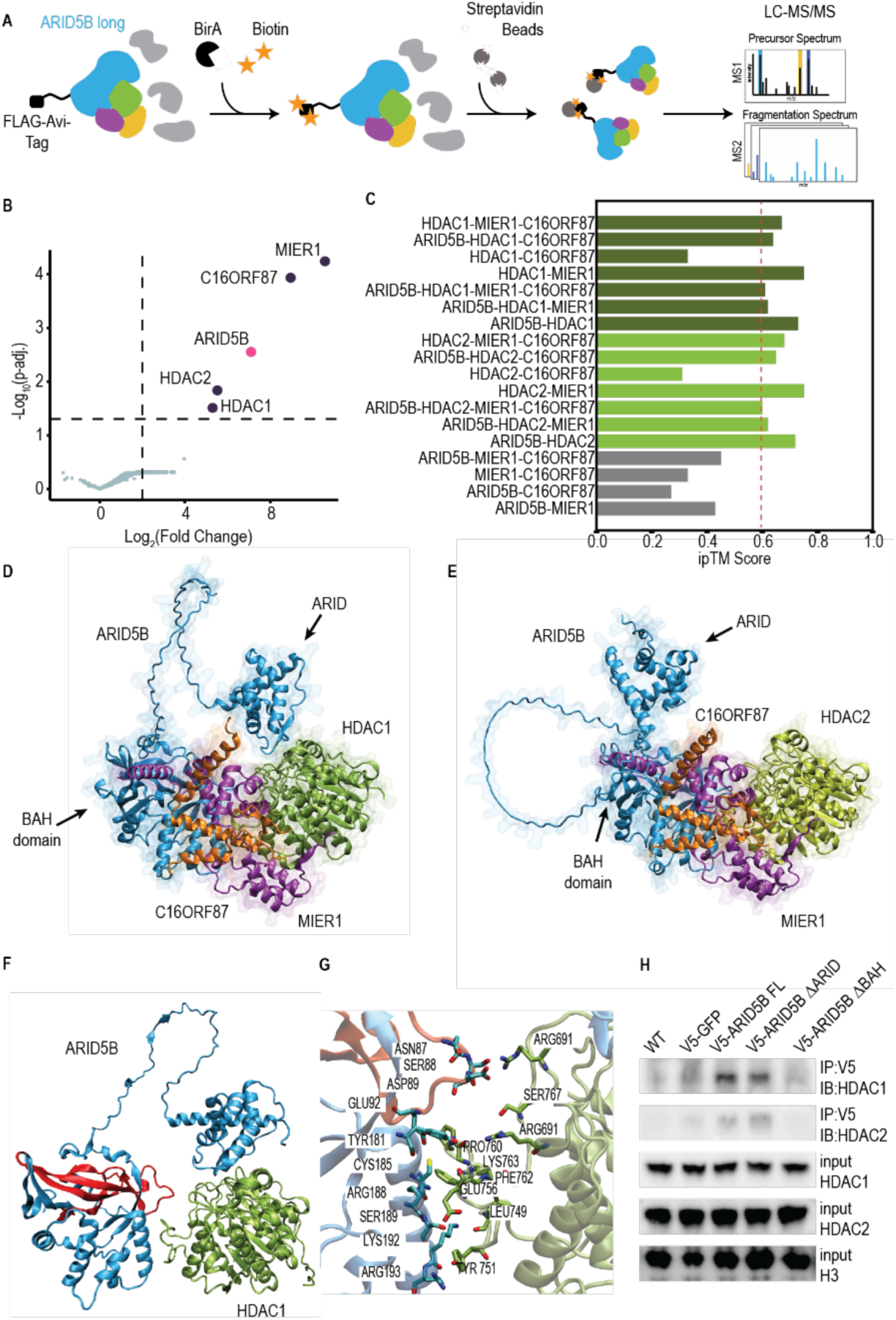
ARID5B forms a protein complex with HDAC1, HDAC2, MIER1 and C16ORF87. **a**. Experimental overview of the identification of ARID5B interaction partners. A construct encoding the murine ARID5B containing an N-term FLAG-AviTag was lentivirally overexpressed in HEK293T cells expressing the Biotin ligase BirA. The latter biotinylates the AviTag in the presence of Biotin, found in the cell culture medium. ARID5B and its interactors were then pulled down using streptavidin beads and quantified by mass spectrometry. **b**. Scatterplot showing enriched proteins in the AP-MS of ARID5B against the Avi-Tag negative control (n=3 biological replicates for samples and controls). Dotted lines indicate a log2FC threshold of larger or equal to 2 and adjusted p-value threshold of smaller or equal to 0.05 (linear model followed by eBayes for moderated t-statistics, and p-value adjustment with Benjamini-Hochberg procedure). Scored interaction partners (purple) and bait (pink) are labeled. **c**. ipTM scores from AlphaFold3 models for all predicted complex combinations. **d., e**. AlphaFold3-predicted structures of ARID5B-MIER1-C16ORF87 in complex with either HDAC1 **(d)** or HDAC2 **(e)**. ARID5B is shown in blue, MIER1 in purple, C16ORF87 in orange, and HDAC1 **(d)** or HDAC2 **(e)** in green/light green, respectively. Disordered regions with low pLDDT scores are omitted for clarity. **f**. ARID5B (blue) in complex with HDAC1 (green) as sampled during molecular dynamics simulations. The BAH domain of ARID5B is highlighted in red. **g**. Predicted binding interface of ARID5B (blue) and HDAC1 (green), highlighting interacting residues (sticks) identified by GetContacts analysis during molecular dynamics simulations. **h**. Co-immunoprecipitation of V5-tagged GFP, full-length ARID5B (FL) and mutants lacking the ARID (ΔARID) or BAH (ΔBAH) domains followed by immunoblotting of HDAC1 and HDAC2 in HAP1 cells.

To better understand the organization of ARID5B with its interactors, we performed structural predictions of complexes containing different interaction partners (**Fig. 1C-E**; **Fig. S1D, Table S3**). Complexes containing ARID5B, MIER1, C16ORF87 and HDAC1 or HDAC2 displayed interface predicted template modelling (ipTM) scores above 0.6, strongly supporting the assembly of ARID5B into a complex (**Fig. 1C**). Interestingly, the prediction of complexes containing C16ORF87 and HDAC1 or HDAC2 drastically improved upon addition of MIER1 and complexes lacking HDAC1 and HDAC2 had the lowest ipTM scores. Chain-pair ipTM scores further predict strong interactions of HDAC1 and/or HDAC2 with all subunits (**Fig. S1E-F**). Notably, among the predicted complex-subunit interactions, ARID5B exhibits the highest-confidence score with HDAC1 and HDAC2, as indicated by an ipTM interaction score of 0.73 and 0.72, respectively (**Fig. S1E-F**). Together with the overall complex structure, this suggests that ARID5B and HDAC1/2 form a scaffold, which is further supported by interactions of MIER1 and C16ORF87 with both ARID5B and HDAC1 or HDAC2.

While ARID5B consists mainly of intrinsically disordered regions, the BAH (aa 34–108) and ARID (aa 319– 411) domains at the N-terminus, have an average pLDDT score of 82, indicating a more confidently predicted and well-defined structure **(Fig. S1G-H**). The N-terminus of ARID5B is also predicted to interact with HDAC1 and HDAC2, with residues 1-250 consistently having the lowest prediction alignment error across models with either HDAC1 or HDAC2 (**Fig. S1I-J**). Molecular dynamics simulations of the ARID5B– HDAC1 complex revealed that residues Asn87, Ser88, Asp89, and Glu92 of the BAH domain, along with an α-helix of ARID5B (aa 181–193), are located at the interface and contribute to stabilizing the interaction with HDAC1 (**Fig. 1F-G**). Co-immunoprecipitation of murine V5-tagged ARID5B, and domain deletion mutants (ΔARID and ΔBAH) in HAP1 cells confirmed that the interaction of ARID5B with HDAC1 and HDAC2 is mediated by the BAH domain (**Fig. 1H**; **Fig. S1K-L**). This further suggests that while the BAH domain of ARID5B is responsible for protein-protein interactions in the complex, ARID may remain accessible for DNA binding.

### ARID5B localizes on active genomic regions

To map ARID5B binding on the genome, we performed CUT&RUN in wild-type (WT) and ARID5B KO NALM6 cells, a cell line derived from a B-ALL patient (**Fig. S2A-C**). We identified 4,200 genomic regions occupied by ARID5B in NALM6 cells (**Fig. S2D**; **Table S4**). Loci occupied by ARID5B enriched for motif and binding of B cell lineage-specific transcription factors, such as IKZF1, EBF1, PAX5, and TCF3 (**Fig. 2A-B**). ARID5B co-localized with the active histone marks H3K27ac and H3K4me1 and did not co-localize with the repressive histone mark, H3K9me2, the substrate of PHF2^26,27^ (**Fig. 2C-D**). Chromatin state analysis by ChromHMM indicated that ARID5B localizes in active TSSs and enhancers (**Fig. 2E**), with 9.8% and 12.7% of ARID5B localizing to TSS proximal and distal and 29.7% and 35% of ARID5B peaks occupying intronic and intergenic genomic regions, respectively (**Fig. 2F**). To observe the consequences of ARID5B loss globally on chromatin, we extended our CUT&RUN analysis to NALM6 KO cells and compared the distribution of H3K4me1 and H3K27ac in WT and ARID5B KO NALM6 cells. Importantly, 980 loci gained H3K27ac and 428 loci lost it upon ARID5B KO, while 976 and 852 regions gained and lost H3K4me1 in KO cells, respectively (**Fig 2G-H**, **Fig. S2E, Table S5**). In fact, H3K27ac signal intensity was markedly higher on ARID5B peaks in KO compared to WT cells (**Fig. 2I, Fig. S2F**). Together, this suggests that ARID5B modulates H3K27ac levels in active genomic regions.

**Figure 2:**
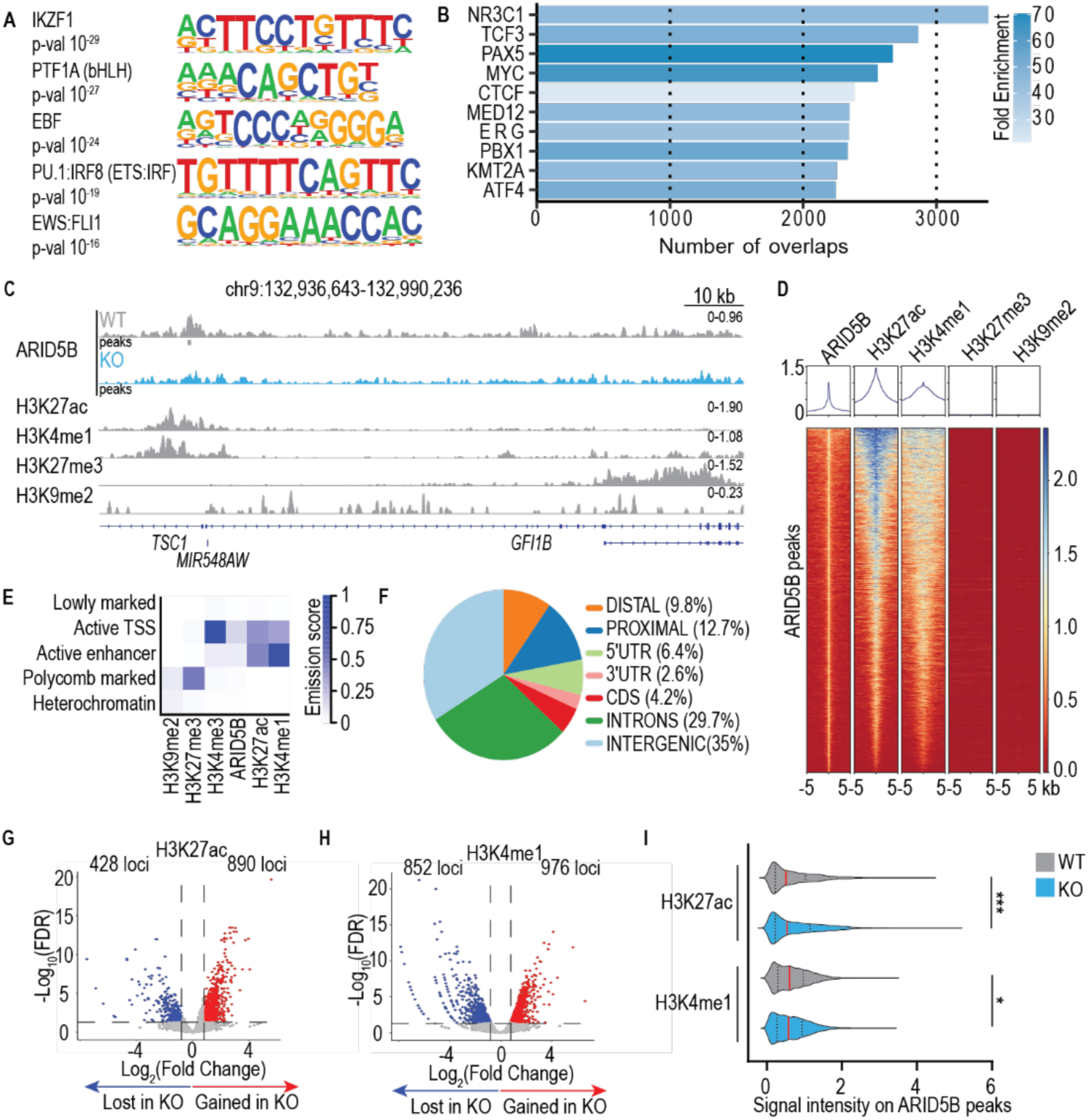
ARID5B localizes on active genomic regions. **a**. Five most significant HOMER *de novo* motifs enriched on 4,200 ARID5B peaks. **b**. Enrichment analysis using ChIP Atlas of publicly available TFs or co-factors co-localizing with 4,200 loci occupied by ARID5B in NALM6 WT cells. The analysis was restricted to factors probed in blood cells by setting the “Cell Class” filter to “Blood”. The bar plot depicts the top factors and their overlap with ARID5B occupied regions. Only data of endogenous, non-tagged, ChIP-seq in “blood” cells (depicted are overlaps of ChIP-seq peaks in NALM6, RCH-ACV, SEM, Acute Lymphoid Leukemia, RS-411, Pre-B cells, OCI-LY1) with an FDR ≤ to 0.05 is shown. **c.** Genome-wide coverage of ARID5B in WT and ARID5B KO NALM6 cells, and of active (H3K27ac, H3K4me1) and repressive (H3K27me3, H3K9me2) histone marks in NALM6 WT cells at the *FAM1638* locus. The coverage of two merged biological replicates per histone mark and of ARID5B is shown. **d.** Aggregate plots and heatmaps of ARID5B, active and repressive histone marks in NALM6 cells centered on the previously determined 4,200 ARID5B peaks. **e.** ChromHMM-defined chromatin states in NALM6 cells based on ARID5B occupancy and active (H3K27ac, H3K4me1, H3K4me3) and repressive (H3K27me3, H3K9me2) histone marks. The coverage of two merged biological replicates per histone mark was used for the analysis. **f.** Genomic distribution of the 4,200 ARID5B peaks called in NALM6 cells. **g., h.** Volcano plots of H3K27ac **(g)** and H3K4me1 **(h)** differentially bound sites upon ARID5B loss. EdgeR-based DiffBind analysis was used to identify loci losing (blue) or gaining (red) H3K27ac or H3K4me1 occupancy in ARID5B KO cells compared to NALM6 WT counterparts. Dotted lines indicate a log2FC threshold of larger or equal to 0.8 and a FDR threshold of smaller or equal to 0.05. Two biological replicates per histone mark per genotype were used for the analysis.**i.** Violin plot of H3K27ac and H3K4me1 average signal intensity on 4,200 ARID5B peaks in WT and ARID5B KO NALM6 cells. The average signal intensity for each region was calculated within +/-5 kb bins using *compute matrix* from deeptools. The median signal intensity is shown in red and quartiles are depicted by dotted lines. For statistics, the nonparametric and un-paired Kolmogorov-Smirnov test was performed. The coverage of two merged biological replicates per histone mark per genotype was used for the analysis. \**P* ≤ 0.05, \*\**P* ≤ 0.01, \*\*\**P* ≤ 0.001, ns = not significant.

### ARID5B recruits HDAC1 and HDAC2 to the genome

The fact that ARID5B interacts with histone deacetylases and that inactivation of ARID5B leads to a gain in H3K27ac, suggests a loss of HDAC activity at ARID5B occupied sites. Thus, to further dissect the chromatin regulatory role of the ARID5B complex, we performed CUT&RUN of HDAC1 and HDAC2. Strikingly, we observed that 76.3% (3,206 peaks) and 77.4% (3,250 peaks) of ARID5B bound sites are occupied by HDAC1 and HDAC2, respectively, with 2,872 ARID5B peaks being bound by all three factors (**Fig. 3A**; **Table S6**). To further investigate the interaction of ARID5B with HDACs and the assembly of the complex on chromatin, we compared HDAC binding in WT and ARID5B KO cells. We uncovered that HDAC1 and HDAC2 occupancy on ARID5B peaks was reduced upon loss of ARID5B (**Fig. 3B-C**). Strikingly, 12,765 and 6,061 sites significantly lost HDAC1 and HDAC2 signal upon ARID5B loss, respectively (**Fig. 3D-F**; **Table S7**). Importantly, no changes in *HDAC1* and *HDAC2* mRNA levels were observed upon ARID5B KO (**Fig. 3G**), indicating that ARID5B does not regulate HDAC1 and HDAC2 expression but rather tethers these factors to chromatin.

**Figure 3:**
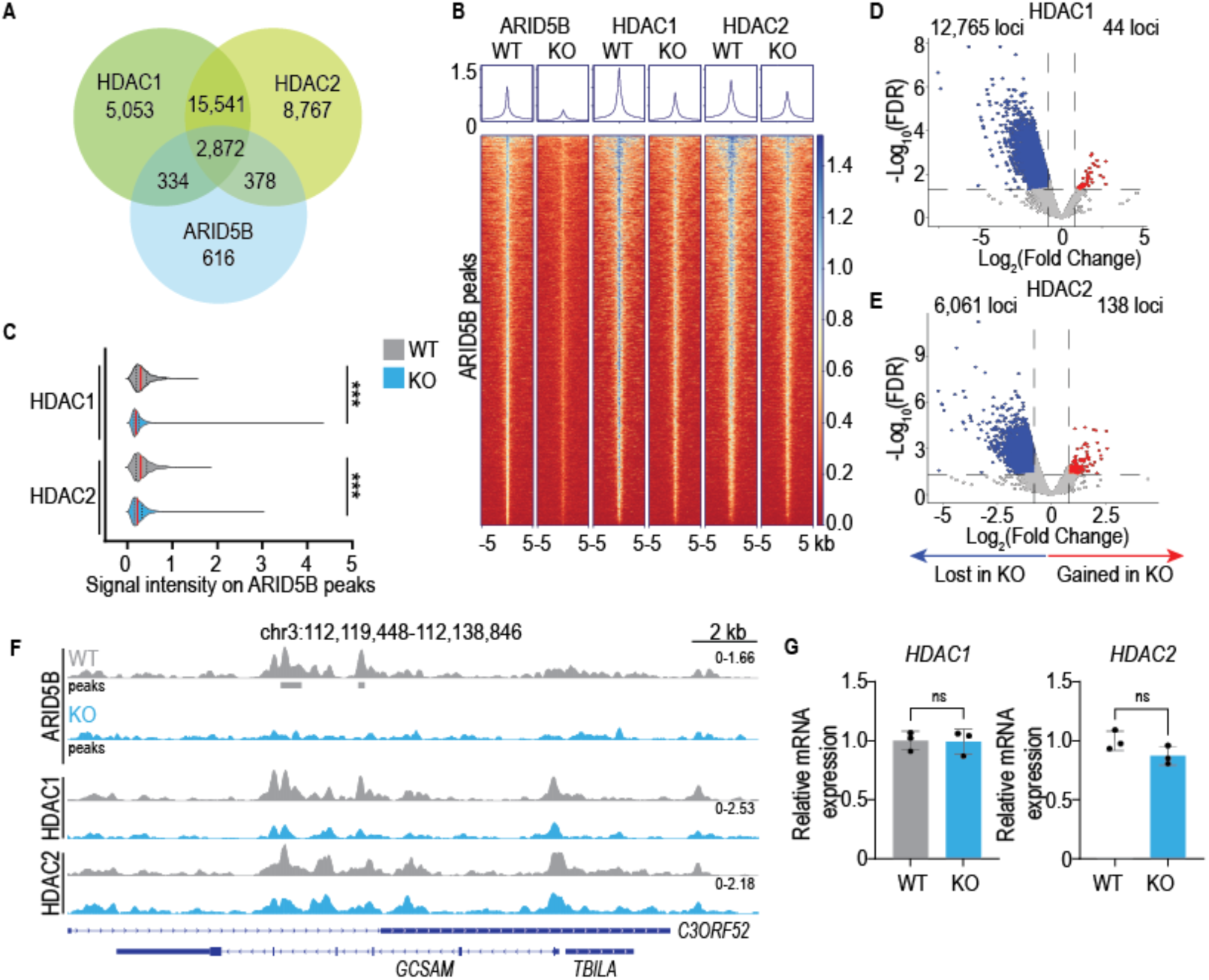
ARID5B tethers HDAC1 and HDAC2 to the genome. **a**. Venn Diagram of ARID5B, HDAC1 and HDAC2 peaks in NALM6 WT cells. **b.** Aggregate plots and heatmaps of ARID5B, HDAC1 and HDAC2 in WT and ARID5B KO NALM6 cells centered on the previously determined 4,200 ARID5B peaks. The coverage of two merged biological replicates per histone mark per genotype was used for the analysis. **c.** Violin plot of HDAC1 and HDAC2 average signal intensity on 4,200 ARID5B peaks in WT and ARID5B KO NALM6 cells. The average signal intensity for each region was calculated within +/-5kb bins using *compute matrix* from deeptools. The median signal intensity is shown in red and quartiles are depicted by dotted lines. For statistics, the nonparametric and un-paired Kolmogorov-Smirnov test was performed. The coverage of two merged biological replicates per histone mark per genotype was used for the analysis. \**P* ≤ 0.05, \*\**P* ≤ 0.01, \*\*\**P* ≤ 0.001, ns = not significant. **d., e.** Volcano plots of HDAC1 **(d)** and HDAC2 **(e)** differentially bound peaks upon ARID5B loss. EdgeR-based DiffBind analysis was used to identify loci losing (blue) or gaining (red) HDAC1 or HDAC2 occupancy in ARID5B KO compared to WT NALM6 cells. Dotted lines indicate a log2FC threshold of larger or equal to 0.8 and a FDR threshold of smaller or equal to 0.05. Two biological replicates for HDAC1 or HDAC2 per genotype were used for the analysis. **f.** Genome-wide coverage of ARID5B, HDAC1 and HDAC2 in WT and ARID5B KO NALM6 cells at the *POU2F2* locus. The coverage of two merged biological replicates of HDAC1, HDAC2 or ARID5B per genotype is shown. **g.** *HDAC1* and *HDAC2* expression in NALM6 WT and ARID5B KO cells assessed by RT-qPCR. Relative expression was obtained upon normalization to the housekeeping gene, *GUSB*. Individual data points, mean and standard deviation are depicted by dots, bars and whiskers, respectively. For statistics, unpaired two-tailed student’s t-test was run with *n* = 3 and \**P* ≤ 0.05, \*\**P* ≤ 0.01, \*\*\**P* ≤ 0.001, ns = not significant. The qPCR was run using technical duplicates of every biological replicate (*n* = 3) sample.

### Loss of ARID5B leads to de-repression of HDAC target genes altering B cell-specific programs

To assess whether ARID5B mediates the repression of those sites showing loss of HDAC1 and HDAC2, we conducted H3K27ac CUT&RUN upon Entinostat treatment, a HDAC Class I inhibitor^28^ (**Fig. 4A**; **Table S8**). As HDAC1 and HDAC2 catalyze the de-acetylation of H3K27ac^29,30^, we compared peaks with gain of H3K27ac upon HDAC inhibition, with peaks losing HDAC1 or HDAC2 upon ARID5B KO. Indeed, we observe that 37.6% (553 peaks) and 21.3% (313 peaks) of regions gaining H3K27ac upon Entinostat treatment lose HDAC1 or HDAC2 binding upon ARID5B loss (**Fig. 4B, C**; **Table S8**). Together, this suggests that a subset of HDAC-regulated loci are regulated by ARID5B in concert with HDAC1 and/or HDAC2, forming a chromatin repressor complex.

**Figure 4.**
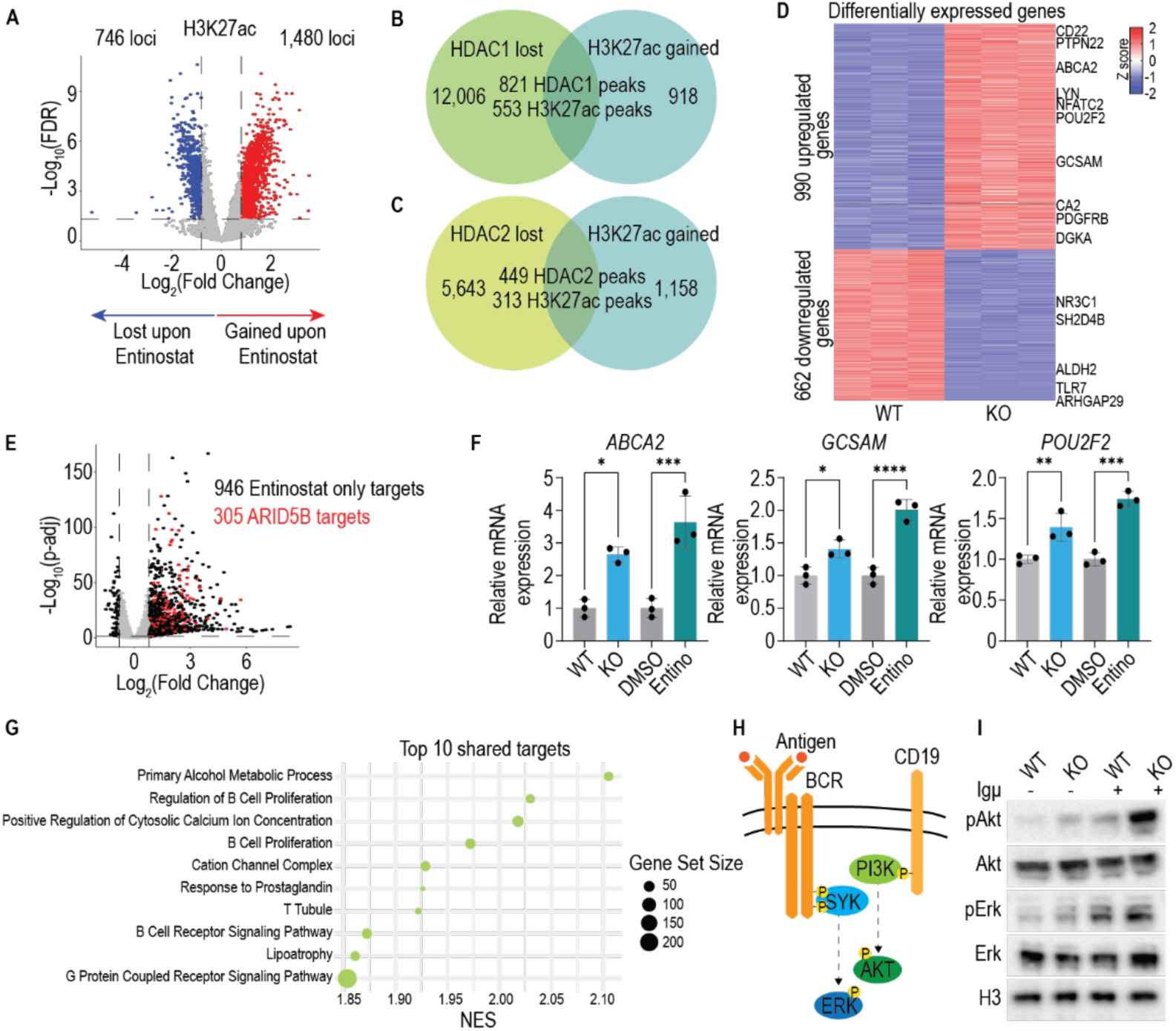
ARID5B and HDACs regulate B-cell specific processes. **a**. Volcano plot depicting differential occupancy of H3K27ac upon Entinostat (MS-274) treatment in NALM6 WT cells. EdgeR-based DiffBind analysis was used to identify loci losing (blue) or gaining (red) H3K27ac occupancy in Entinostat (MS-275) treated cells compared to vehicle-treated NALM6 WT counterparts. Dotted lines indicate a log2FC threshold of larger or equal to 0.8 and a FDR threshold of smaller or equal to 0.05. NALM6 WT cells were treated with vehicle or 500 nM Entinostat (MS-275) for 24 h prior to CUT&RUN experiment. Two biological replicates per treatment were used for the analysis. **b., c.** Venn Diagram of peaks losing HDAC1 **(b)** or HDAC2 **(c)** occupancy upon ARID5B loss and peaks gaining H3K27ac upon HDAC inhibition by Entinostat in NALM6 WT cells. Differentially bound sites were identified using the DiffBind (edgeR mode) package and the following threshold were set for lost (log2FC ≤ -0.8, FDR ≤ 0.05) and gained regions (log2FC ≥ 0.8, FDR ≤ 0.05). NALM6 WT cells were treated with vehicle or 500 nM Entinostat (MS-275) for 24 h prior to CUT&RUN experiment. Two biological replicates per treatment and per HDAC1 or HDAC2 per genotype were used for the analysis. **d.** Heatmap of up- and downregulated genes upon ARID5B loss. Depicted are the Z scores for each regulated gene across biological triplicates in NALM6 WT and ARID5B KO cells. The following thresholds were used for up (log2FC ≥ 0.8, padj ≤ 0.05, cpm (KO) ≥ 30) and downregulated (log2FC ≤ -0.8, padj ≤ 0.05, cpm (WT) ≥ 30) genes. Representative genes are shown. **e.** Volcano plot of genes regulated upon Entinostat (MS-275) treatment in NALM6 WT cells. Highlighted in red are Entinostat-responsive genes that are also upregulated in ARID5B KO cells. Dotted lines indicate a log2FC threshold of larger or equal to 0.8 and a FDR threshold of smaller or equal to 0.05. The following thresholds were used for up (log2FC ≥ 0.8, padj ≤ 0.05, cpm (KO/Entinostat) ≥ 30) and downregulated (log2FC ≤ -0.8, padj ≤ 0.05, cpm (WT/DMSO) ≥ 30) genes. NALM6 WT cells were treated with vehicle or 500 nM Entinostat (MS-275) for 24 h prior to mRNAseq experiment. Biological triplicates per genotype and per treatment were used. **f.** Expression of ARID5B and Entinostat target genes. Relative expression was obtained upon normalization to the housekeeping gene, *GUSB*. Individual data points, mean and standard deviation are depicted by dots, bars and whiskers, respectively. For statistics, one-way ANOVA followed by Tukey multiple comparison test was run with *n* = 3 and \**P* ≤ 0.05, \*\**P* ≤ 0.01, \*\*\**P* ≤ 0.001, ns = not significant. The qPCR was run using technical duplicates of every biological replicate (*n* = 3) sample. **g.** Top 10 GO-terms enriched in gene set enrichment analysis (GSEA – C2 collection) of ARID5B KO and Entinostat treated NALM6 cells. Depicted are the normalized enrichment scores (NES) and the number of genes contained in each gene ontology term. GSEA was run comparing NALM6 WT with ARID5B KO cells and comparing vehicle-treated with Entinostat (MS-275)-treated cells. Terms with an FDR smaller or equal to 0.25 were considered statistically significant. NALM6 WT cells were treated with vehicle or 500 nM Entinostat (MS-275) for 24 h prior to mRNAseq experiment. Biological triplicates per genotype and per treatment were used. **h.** Simplified schematic of B cell receptor (BCR) and the co-receptor CD19, as well as downstream signaling intermediates, which are activated upon antigen binding to BCR. **i.** Western Blot depicting pAkt, Akt, pErk, Erk levels in NALM6 WT and ARID5B KO cells stimulated with 10 µ/ml Igµ for 10 min.

To investigate the effect of the ARID5B complex on transcription, we performed mRNAseq on WT and ARID5B KO cells. This analysis revealed that more genes are de-repressed and thus more highly expressed upon ARID5B loss, with 990 upregulated and 662 downregulated genes (**Fig. 4D**; **Fig S3A**; **Table S9**). Interestingly, 305 genes are upregulated upon ARID5B loss and HDAC inhibition by Entinostat treatment, suggesting that these genes are regulated by the ARID5B repressor complex in a HDAC-dependent manner (**Fig. 4E-F**; **Table S9**). Gene ontology analysis showed that genes upregulated only upon ARID5B loss or only upon HDAC inhibition are mainly involved in DNA replication or antigen presentation, respectively (**Fig. S3B-C**; **Table S10**). On the other hand, genes upregulated in KO and upon Entinostat treatment, thus repressed by the ARID5B complex, are implicated in B-cell specific cellular processes and signaling pathways (**Fig. 4G**; **Table S10**).

Importantly, targets of the ARID5B repressor complex are involved in B cell proliferation, calcium signaling regulation and BCR signaling, suggesting that the ARID5B repressor complex plays a crucial role in BCR signaling regulation. To test this, we stimulated WT and ARID5B KO cells with Igµ and observed that both showed increased Erk phosphorylation upon stimulation **(Fig. 4H-I**). Interestingly, KO cells displayed an additional increase in phosphorylated Akt levels upon stimulation, suggesting a difference in antigen-dependent signaling (**Fig. 4H-I**).

Taken together, we show that ARID5B associates with HDAC1, HDAC2, MIER1 and C16ORF87 forming a novel co-repressor complex. ARID5B further tethers the complex to TSSs and distal regulatory elements, actively repressing B cell-specific genes (**Fig. 5**).

**Figure 5:**
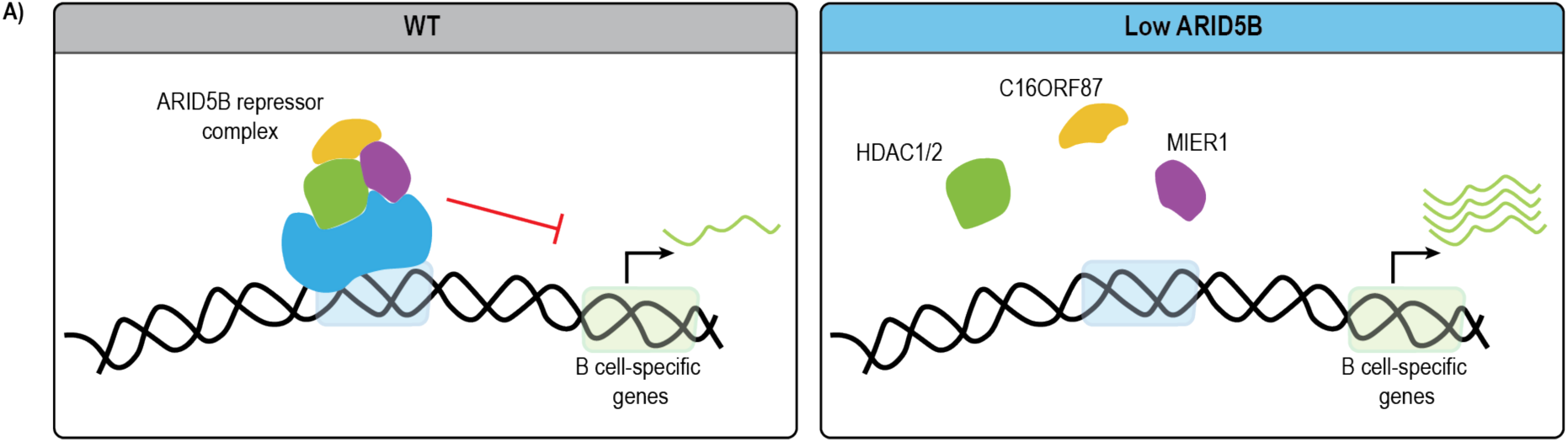
The ARID5B repressor complex regulates the expression of B cell specific genes. **a.** Schematic depicting the role of the ARID5B repressor complex on chromatin. ARID5B interacts with HDAC1 or HDAC2 (green), MIER1 (purple) and C16ORF87 (orange) and forms a repressor complex. It further tethers HDAC to promoter proximal and distal regions this way repressing genes involved in B cell proliferation and B cell-specific signaling. In the absence of ARID5B, HDAC occupancy at those sites is reduced and consequent target gene expression is increased.

## Discussion

Here we show that ARID5B forms a repressor complex with HDAC1, HDAC2, MIER1 and C16ORF87. Importantly, the BAH domain of ARID5B mediates the interaction with HDAC1 and HDAC2, which is further supported by MIER1 and C16ORF87 binding. On a genome-wide level, ARID5B co-localizes with active histone marks, occupying mainly intergenic and intronic regions, while also binding TSS proximal and distal loci. ARID5B tethers HDAC1 and HDAC2 to chromatin, and loss of ARID5B leads to decreased HDAC occupancy, resulting in upregulation of a subset of target genes. Together, ARID5B and HDACs regulate B cell specific genes, fine-tuning BCR signaling responses.

One could speculate that the ARID5B repressor complex activity changes during B cell development and is modulated by B cell-specific signaling. Importantly, ARID5B expression continuously increases from the pro-B to large and small pre-B cell stages^16^, along with the maturation of BCR^31,32^. This suggests that the ARID5B repressor complex is a crucial player in regulating pre-BCR and BCR signaling during development. Interestingly, a BCR signaling second messenger, Inositol 1,4,5-triphosphate (InsP_3_) ^31,33^, together with its phosphorylated version, Inositol 1,4,5,6-tetraphosphate, have been shown to enhance the binding of ELM2-SANT domain proteins, such as MIER1, to HDAC and this way promote HDAC activity^34,35^. Thus, one could speculate that not only the expression of ARID5B during B cell development, but also onset pre-BCR signaling and changes in cellular InsP_3_ levels, regulate and serve as feedback loops to the timely activity of the ARID5B repressor complex.

Interestingly, the expression of both ARID5B isoforms is differently associated with B-ALL outcome. While increased expression of ARID5B short isoform is associated with lower overall and event-free survival, higher levels of the ARID5B long isoform (that retains the BAH domain required to interact with HDACs), are associated with increased event-free survival^36^. This suggests that the long isoform of ARID5B plays a rather tumor suppressive role, implying that the presence of a BAH domain and consequently the activity of the ARID5B repressor complex is crucial to control transcripts associated with oncogenesis. Alternatively, the short isoform of ARID5B could exert a dominant negative function, which when overexpressed, outcompetes the ARID5B repressor complex on chromatin, thus de-repressing B cell specific genes.

Finally, altered activity of HDAC-containing repressor complexes, such as NCoR and NuRD, has been shown to promote the development of hematological malignancies, such as lymphomas^37,38^ and leukemias^39^, respectively. While NCoR interacts with ERG to control leukemia-associated genes^39^, loss of NuRD favors T cell lymphoma by de-repressing E2A and creating a B and T cell progenitor imbalance^37^. In fact, Mbd3/NuRD ablation in hematopoietic stem cells leads to premature and increased open chromatin regions at B cell lineages specific enhancers and consequent faster commitment to the B cell lineage^37^. Still, NuRD loss leads to decreased B cells at early developmental stages, affecting especially the pro-B cell stage and the ratio of large to small pre-B cells^40^. The latter is due to the dysregulation of cell cycle genes and consequent cell cycle arrest of large pre-B cells^41^. Similarly, HDAC1 and HDAC2 have been shown to regulate cell cycle in pre-B and proliferating mature B cells^42^. Thus, HDAC and its complexes have been described to control a plethora of B cell specific processes pre-disposing to leukemia.

Future work will be necessary to study the contribution of the ARID5B repressor complex to B cell development *in vivo*. Of importance, is to distinguish the contribution of HDAC1 and HDAC2 in the context of ARID5B as well as other repressor complexes. Monitoring and characterizing the onset of leukemia in mice depleted from components of the ARID5B repressor complex will further provide insights into the contribution and role of the complex in leukemia predisposition. Together, this will elucidate the molecular mechanism by which ARID5B pre-disposes to B-ALL, providing new insights into potential therapeutic dependencies, such as HAT inhibitors.

## Resources

### Lead contact

Further information and requests for resources and reagents should be directed to and will be fulfilled by the lead contact.

## Materials availability

Cell lines and expression vectors generated in this study are available upon request.

## Data and code availability

The next-generation sequencing data discussed in this publication have been deposited in NCBI’s Gene Expression Omnibus^43^ and will be accessible through GEO Series accession number GSE297401 after peer review. Publicly available datasets used in this study are accessible at GSM1080936. Scripts used in this study to analyze CUT&RUN and mRNAseq datasets have been deposited on Github (https://github.com/seruggialab).

The mass spectrometry proteomics data have been deposited to the ProteomeXchange Consortium via the PRIDE^44^ partner repository with the dataset identifier PXD061867 and will be accessible after peer review. Predicted structures of protein interactions identified in the ARID5B co-IP are available at https://zenodo.org/records/17343031.

## Acknowledgements

This research was funded in whole or in part by the Austrian Science Fund (FWF) [10.55776/P36302]. For open access purposes, the author has applied a CC BY public copyright license to any author accepted manuscript version arising from this submission. Research in the Seruggia laboratory is further supported by the WES Leukemia Research Foundation, the Austrian Science Fund (FWF, project P36069), the European Research Council (ERC) under the Horizon 2020 research and innovation programme (grant agreement 947803) and under Marie Skłodowska–Curie (grant agreement 101061151). G.S.-F. is supported by the Austrian Academy of Sciences. The authors thank the Flow Cytometry Facility at CCRI and the Biomedical Sequencing Facility at CeMM. We thank the Proteomics team of the Molecular Discovery Platform at CeMM for mass-spectrometric data acquisition and want to especially acknowledge Thomas Hannich and Anna Tinnacher. We thank Georg Winter (CeMM, Vienna) for the pLEX305 N-term miniturbo V5 plasmid. The authors also thank Zain Patel (MGH, Boston) and Animesh Awasthi (CeMM, Vienna) for fruitful discussions on data analysis.

## Author Contributions

Conceptualization: A.P.K. and D.S.; Formal analysis: A.P.K, F.F., B.S.; Funding acquisition: A.P.K. and D.S.; Investigation: A.P.K, F.F., B.S., S.M., P.B., G.K.; Methodology: A.P.K, F.F., B.S.; Resources: D.S.; Supervision: D.S., G.S.F; Writing - original draft: A.P.K., D.S.; Writing – review and editing: A.P.K., F.F., B.S., D.S.

## Supplemental information

**Figures S1–S3**

**Table S1. Experimentally mapped protein interactions of ARID5B.**

**Table S2. Reported PPIs between subunits of ARID5B repressor complex.**

**Table S3. Summary of top AlphaFold3-predicted protein complex models.**

**Table S4. ARID5B peaks in NALM6 WT and ARID5B KO cells.**

**Table S5. Differential binding of H3K27ac and H3K4me1 in NALM6 WT and ARID5B KO cells.**

**Table S6. Regions occupied by HDAC1, HDAC2 or ARID5B, HDAC1 and HDAC2 in NALM6 WT cells.**

**Table S7. Differential binding of HDAC1 and HDAC2 in NALM6 WT and ARID5B KO cells.**

**Table S8. Regions gaining H3K27ac in NALM6 WT cells upon Entinostat treatment and losing HDAC1 or HDAC2 binding upon ARID5B KO.**

**Table S9. Genes regulated by ARID5B and/or upon Entinostat treatment.**

**Table S10. GO terms enriched in NALM6 WT vs ARID5B KO and/or DMSO vs Entinostat treatment. Table S11. Oligonucleotides, plasmids and antibodies used in this study**

## Materials and Methods

### Cell culture

NALM6 (CVCL_0092), HEK293T (CVCL_0063) and HAP1 (CVCL_Y019) were cultured in RPMI 1640 (Gibco, Life Technologies), DMEM (Gibco, Life Technologies) and IMDM (Gibco, Life Technologies), respectively, all supplemented with 10% Fetal Bovine Serum, 100 U/mL Penicillin-Streptomycin and 2 mM L-Glutamine at 37 °C, 5% CO2. Cells were tested weekly for mycoplasma contamination (MycoAlert, Lonza LT07-318).

### Generation of NALM6 KO cells

For the KO of *ARID5B,* Cas9-stably expressing NALM6 cells were electroporated with RNP complexes containing sgRNA targeting exons 5 and 10 of the long isoform of *ARID5B* (Synthego, Table S11 - Oligonucleotides). Cells were directly seeded in 96 well plates at a density of 5 cells/well to obtain clones. KO clones were identified by genotyping followed by Sanger sequencing (see Table S11 – Oligonucleotides).

### Genotyping of KO cells

For the validation of KO cells, the region targeted by the sgRNA was amplified by PCR (see Table S11 – Oligonucleotides) and editing was assessed by sanger sequencing. Briefly, cells were lysed in Pawell’s buffer (10 mM Tris-HCl (pH 9.0), 50 mM KCl, 0.45% NP-40, 0.45% Tween-20) with 0.8 mg/mL Proteinase K (Qiagen, #19133) at 56 °C for 15 min followed by 95 °C for 5 min. OneTaq DNA Polymerase (NEB, #M0486) was used for PCR with 1.5 µL of DNA as input.

### Treatment of NALM6 WT with HDAC inhibitors

Inhibition of HDAC Class I was achieved upon treating NALM6 Cas9 or ARID5B KO cells at a density of 1 million cells/mL with 500 nM Entinostat – MS-275 (Selleckchem, #S1053) or 0.005% DMSO for 24 h prior to RNA harvesting or CUT&RUN experiment.

### Stimulation of NALM6 WT and ARID5B KO cells with Igµ

NALM6 WT and ARID5B KO cells were seeded at a density of 2 million cells/mL and stimulated with 10 µg/mL Goat F(ab’)2 Anti-Human IgM-UNLB (Southern Biotech, #2022-01) for 10 min prior to protein harvesting.

### Protein Harvesting

Cells were lysed using NP-40 buffer (50 mM Tris-Hcl (pH 8.0), 150 mM NaCl, 1% NP-40) supplemented with 1:100 Protease Inhibitor Cocktail (Sigma, #P8340) and 0.2 mM Phenylmethanesulfonyl fluoride (Sigma, #93482). Lysates were incubated for 15 min at 4 °C and centrifuged for 10 min at top speed at 4 °C. The supernatant was transferred to a fresh tube and samples were prepared for Western Blot.

### Lentivirus production

HEK293T cells were seeded in 15 cm plates the day prior to transfection. When 80% confluent, cells were transfected with 8.75 µg VSV-G, 16.25 µg, pPAX2, and 25 µg of the lentivector of interest (see Table S11 – Plasmids) using 150 µg branched PEI (Sigma, #408727). The following day, the media of the cells was changed with fresh media. Lentiviral supernatant was collected 2 and 3 days after transfection, centrifuged at 2,500 rpm for 5 min, filtered using a 0.45 µm filter (Thermo Scientific, #723-2545) and concentrated by ultracentrifugation at 24,000 rpm, at 4 °C for 2 h, using 20% sucrose in PBS. For crude lentivirus preparation, transfected HEK293T cells were cultured in either DMEM, IMDM or RPMI, the lentivirus supernatant was collected 2 and 3 days post-transfected, centrifuged at 2,500 rpm for 5 min and filtered using a 0.45 µm filter (Thermo Scientific, #723-2545).

### Lentivirus transduction

Cells were seeded at a density of 1 million cells/mL and 10 µL of purified lentivirus or an equal volume of crude lentivirus was added. The day after transduction, 10 µg/mL of Blasticidin (Gibco, #A1113903) or 1 µg/mL of Puromycin (Thermo Fisher, #A1113803) were used to select cells transduced with viruses carrying a Blasticidin or Puromycin resistance cassette, respectively. Cells transduced with constructs containing a GFP cassette were FACS sorted for GFP 3-5 days post-transduction.

### Affinity purification of FLAG-Avi-tagged ARID5B followed by Mass Spectrometry (AP-MS)

The murine *Arid5b* ORF with 5’ N-terminus FLAG-AviTag was synthesized (IDT) and cloned into a lentivector under the EF1a promoter using Gibson Assembly Master Mix (NEB, #E2611S) (see Table S11 – Plasmids). HEK293T cells constitutively expressing the E.Coli Biotin ligase BirA were transduced with the ARID5B lentivector and selected with Puromycin for 5 days. One-step affinity purification with streptavidin beads was performed as previously reported^22,45^. In brief, nuclear extracts were obtained from approximately 300 million cells using NE-PER Nuclear and Cytoplasmic Extraction Reagents (Thermo Fisher Scientific, #78833) and quantified using the Pierce BCA Protein Assay Kit (Thermo Fisher Scientific, #23225). 5 mg of nuclear extract from HEK293T cells either expressing only BirA (control) or BirA plus biotin-tagged ARID5B were diluted in 10 mL of IP350 buffer (20 mM Tris-HCl (pH 7.5), 0.3% NP-40, 1 mM EDTA, 10% glycerol, 350 mM NaCl, 1 mM DTT, 0.2 Phenylmethanesulfonyl fluoride (Sigma, #93482)) supplemented with a Protease Inhibitor Cocktail (Sigma Aldrich, #P8340-5, 1:100). 100 uL Dynabeads MyOne streptavidin T1 beads (Thermo Fisher Scientific, #65601) were added and incubated with rotation overnight at 4 °C. Beads were washed four times with 10 mL of IP350 buffer and eluted in 250 µL Buffer II (50mM HEPES (pH 8.0), 150 mM NaCl, 5 mM EDTA, 2% SDS). Next, eluted samples were diluted to equal amounts in lysis buffer (2% SDS, 50 mM HEPES, pH 8, supplemented with 1 mM Phenylmethanesulfonyl fluoride (Sigma, #93482) and protease inhibitors) up to 100 µL. Next DTT (final concentration of 10 mM DTT) was added, and samples were incubated for 1 h at 56 °C. After reduction, cysteine residues were treated with IAA (final concentration of approximately 50 mM) and incubated in the dark for 30 min at room temperature. To each sample 8 µL washed and equilibrated magnetic SP3 beads (SpeedBeads, GE Healthcare) were added, and proteins were precipitated by the addition of Acetonitrile (final concentration of 70% v/v). The samples were incubated for 18 minutes before the beads were immobilized for 2 min and the supernatant was removed. Beads were rinsed twice with 70% (v/v) ethanol, followed by a wash in acetonitrile (ACN). After the final washing step, 100 µL of 50 mM Ammonium-bicarbonate (NH_4_HCO_3_) was added to the beads, and proteins were digested by the addition of 1.5 µg of sequencing grade trypsin. Samples were incubated on the shaker with low agitation speed overnight (∼ 12 h) at 37 °C. Next, samples were acidified to pH 2-3 with 30% TFA and cleaned up by STAGE-tip C18.

### AP-MS data acquisition

MS-data were acquired on an Orbitrap Fusion Lumos Tribrid mass spectrometer coupled to a Dionex Ultimate 3000 RSLCnano system via a Nanospray Flex Ion Source interface. Peptides were loaded onto the trap column (PepMap 100 C18, 5 µm, 5 × 0.3 mm) and subsequently eluted onto the analytical column (50 cm, 75 µm inner diameter analytical column inhouse packed with ReproSil-Pur 120 C18-AQ, 3 µm, with an ESI emitter (20 µm ID x 7 cm L x 365 µm OD; Orifice ID: 10 µm, CoAnn Technologies) kept at 50 °C. A 120 min analytical gradient with 230 nL/min flow rate was used, using as buffer A 0.4% FA in H2O, and as buffer B 0.4% FA in ACN. The gradient had the following scheme: scheme: 0-4 min 4% buffer B, 4-86 min to 24% B, 86-94 to 36% B, 94-95 min increase to 100% B and 101-102 back to 4% B. The MS was operated in positive mode using a data dependent acquisition (DDA) mode with a 3 sec cycle time. The MS1 scan range was set from 375 – 1650 m/z, and the dynamic exclusion was set to 60 seconds with a mass tolerance of 10 ppm. MS1 spectra were recorded in the Orbitrap at a resolution of 120,000, with a maximum injection time of 80 ms and an AGC target value of 2e5. Only peptide precursor charge stages of 2 to 6 were selected for fragmentation. MS2 fragment spectra were recorded at resolution of 15,000 with an automatic gain control target intensity of 5e4. Fragmentation was achieved with higher energy collision energy (HCD) at 30%. The maximum injection time was limited to 100 ms. The isolation width was set to 1.6 m/z and fragments were recorded from 120 – 2000 m/z.

### MS data processing

MS raw files were processed using FragPipe (version 22.0)^46,47^ and MSFragger (version 4.1)^48^ with the label-free quantification with match-between-runs (LFQ-MBR) workflow. The peptide identification search was conducted against the human protein database downloaded from UniProtKB (consensus sequences, only reviewed, 02/06/2025), which included decoy sequences (reversed protein sequences) and common lab contaminants (e.g., rubber, bovine serum albumin). The default search engine settings of the LFQ-MBR workflow were used for the FragPipe analysis. Briefly, a closed search with the setting “strict trypsin” specificity, N-terminal methionine clipping, and up to two missed cleavages was performed. Up to three variable modifications were allowed, including oxidation on methionine, N-terminal acetylation, and carbamidomethylation of cysteine residues as a static modification. Peptide lengths were set to a minimum of 7 and a maximum of 50 amino acids. MSBooster was enabled, and Percolator was used for PSM validation. ProteinProphet controlled the protein FDR at 0.01. Quantification was performed with IonQuant^47^ using LFQ with MBR enabled, with advanced options set to default.

### AP-MS data analysis

The peptide-level outputs from FragPipe were further analyzed to determine protein abundance and perform differential analysis using R (version 4.4.1). The raw peptide intensities from each MS-injection were normalized based on the total peptide signal and scaled to the median total signal. Peptides which were sparsely identified peptides in less than three samples were excluded from the dataset. Subsequently, the peptides for each protein were summed and ranked across the entire experiment. The six highest abundant peptides (TOP6) were used to infer protein abundances for each sample. If fewer than six peptides were quantified, the protein abundance was inferred from the remaining peptides.

The mean signal of the peptides was used for protein abundance inference. Protein intensities were then log2 transformed, and missing values were imputed by sampling from a normal distribution centered around the lowest 5% quantile of all intensity values. Proteins for which more than 50% of the quantities were imputed were excluded from further analysis. Common lab contaminants (e.g., sheep wool) were also removed. Finally, the log2FC was calculated using the average signal quantified in ARID5B samples compared to the BirA negative controls. To assess enrichment of interaction partners, we applied a linear model to each protein using the limma package^49^. The model estimated the coefficients for each protein, with contrasts between ARID5B and BirA negative controls fitted using empirical Bayes moderation. The resulting statistics were adjusted for multiple testing using the Benjamini-Hochberg method. All proteins with a log2FC larger or equal to 2 and an adjusted p-value of smaller or equal to 0.05 were categorized as interaction partners.

### Reconstruction of literature reported ARID5B PPI-network

PPIs between the ARID5B co-repressor complex (ARID5B, MIER1, C16ORF87, HDAC1 and HADC2) subunits were retrieved from the IntAct^50^ and the BioGRID (version 4.4.243)^51^ databases. Interactions were limited to those identified through affinity enrichment technologies, specifically “Affinity-Capture MS” and “Affinity-Capture Western blot.” Interactions derived from proximity labeling, co-fractionation, and yeast-two-hybrid were excluded. A total of 80 PPIs covering 35 manuscripts were reported among the complex subunits (Table S2). The PPI-network was visualized using Cytoscape (version 3.10.2)^52^.

### Immunofluorescence

Before cell seeding, 8-well Ibidi chambers (Ibidi, 80827) were treated with Poly-D-Lysine (Thermo Fisher, A3890401) diluted 1 in 10 with MilliQ water for 10 min at room temperature, before washing each well 4 times with MilliQ water. For cell seeding, cells were washed once with PBS and incubated with Accutase (Biolegend, 423201) for 3-5 min at room temperature, quenched with complete medium and filtered once through a 35 µm nylon mesh, before cell counting. Cells were seeded at 80,000 cells per well one day prior to fixation in Ibidi chambers. At the time of fixation, cells were washed twice with PBS before fixation with PBS containing 4 % methanol-free formaldehyde (Thermo Fisher Scientific, 28906) for 5 min. The fixative was removed and cells quenched using 10 mM Tris–HCl (Invitrogen, 15567027) pH 7.5 in PBS for 3 min. Cells were permeabilised with PBS containing 0.2 % Triton-X-100 (Sigma Aldrich, X100-100 ml) for 5 min, washed once with PBS to remove residual detergent, and blocked for 30 min with 0.45 µm filtered 2% BSA (Sigma-Aldrich, A9418-50G) in PBS (blocking buffer). Cells were incubated with Streptavidin AF568 (Thermo Fisher, S11225) diluted in blocking buffer for 3 h at room tempeature with gentle shaking, followed by 3 × 10 min washes with PBS containing 1.62 µM Hoechst 33342 (Thermo Fisher Scientific, H3570).

### Microscopy and image processing

Confocal microscopy experiments were imaged on a custom Zeiss LSM 980 microscope fitted with an additional Airyscan2 detector, using a 63× NA 1.4 oil DIC Plan-Aphrochromat (Zeiss) objective and ZEN 3.3 Blue 2020 software.

### Image analysis

Fields of cells were analysed using a custom automated analysis pipeline written in Python. A Gaussian blur was applied to the Hoechst channel using the skimage ‘filters’ module and the Hoechst channel was then segmented using the ‘nuclei’ module of cellpose^53^l. Cells which touched the border of the image were excluded using the ‘clear_border’ functionality from the skimage ‘segmentation’ module. Individual masks were labelled and applied to each cell in the field. The area of the nuclear mask and mean fluorescence within the nuclear mask was then calculated for each segmented cell, and the data output as a csv file for each input condition. Biological replicates were normalised independently, relative to their own internal positive and negative controls and the normalised data then merged for the final figure.

### Structural modeling and molecular dynamics simulations of ARID5B complexes

Structural models of the complexes were predicted using AlphaFold v3.0 (AF3)^54^. Structures were trimmed to remove disordered termini with pLDDT scores below 50. Molecular dynamics (MD) simulations were performed using Amber2024^55^. Systems were prepared with the ff14SB force field^56^ and TIP3P water model^57^. Each complex was solvated in a truncated octahedral box with a 20 Å buffer of TIP3P water and neutralized with Na⁺ and Cl⁻ ions. Energy minimization (40,000 steps) was followed by restrained heating from 0 K to 300 K, NVT equilibration (500 ps), NPT equilibration (500 ps), and a production run of 1 μs. Production simulations were run with pmemd.cuda using periodic boundary conditions, SHAKE^58^ constraints applied to bonds involving hydrogen atoms, and a 4 fs timestep enabled by hydrogen mass repartitioning. Intermolecular contacts were analyzed using GetContacts (https://github.com/getcontacts/getcontacts), and structures were visualized with VMD 1.9.4^59^.

### Co-IP of V5-tagged ARID5B variants

Lentiviral vectors encoding V5-tagged full length murine Arid5b, domain deletions or GFP were obtained by PCR and Gateway cloning (Thermo Fisher Scientific, #11789020 & #11791020) into pLEX305 N-term miniturbo V5 (kind gift from Georg Winter, CeMM). Following lentiviral transduction, HAP1 cells were continuously cultured with Puromycin. Cells from one confluent 15 cm dish were washed with PBS, trypsinized and pelleted at 400 x g for 5 min. For protein harvesting, the cell pellet was resuspended in IP Lysis Buffer (50 mM Tris-HCl (pH 7.5), 150 mM NaCl, 2 mM EDTA, 0.5% Triton X-100, 0.2 mM Phenylmethanesulfonyl fluoride (Sigma, #93482)) supplemented with Benzonase (Millipore Sigma, #706643, 1:200) and Protease Inhibitor Cocktail (Sigma Aldrich, #P8340-%, 1:100). Samples were incubated on ice for 30 min, with in between vortexing every 10 min. Following, the lysate was centrifuged at 10,000 × g for 10 min at 4 °C and the protein in the supernatant quantified using the Pierce BCA Protein Assay Kit (Thermo Fisher Scientific, #23225). V5 Sepharose Beads (Merck, #A7345) were washed five times with IP Lysis Buffer. 1 mg of protein in 400 µL IP Lysis Buffer were incubated with 50 µL of washed V5 Sepharose Beads overnight at 4 °C with rotation. The following day, IP samples were centrifuged at 3,000 × g for 1 min and supernatant (unbound fraction) was collected in a new tube. Beads were washed three times with 500 µL IP Lysis Buffer and proteins eluted from the beads upon addition of 1x Laemmli Sample Buffer (BioRad, #1610747) and incubation at 95 °C for 10 min. Samples were centrifuged at 3,000 × g for 1 min and the supernatant (IP) transferred to a new tube. IP and input samples (18.75 µg - 3.75% of original protein material used for IP) were analyzed by Western Blot.

### Western Blot

Proteins were denatured using XT Sample Buffer (BioRad, #1610791) or 4× Laemmli Sample Buffer (BioRad, #1610747) for 10 min at 95 °C. 4-12% Criterion XT Bis-Tris Protein Gels (BioRad, #3450125) or 4-20% Mini-Protean TGX Precast Protein Gels (BioRad, #4561096) were used to separate the proteins, which were then transferred to low-fluorescence PVDF membranes (BioRad, #1704275) using the High MW program of the Trans-Blot Turbo Transfer System (BioRad, #1704150). The membranes were blocked for 1 h at RT, prior to overnight primary antibody incubation, followed by 2 h RT secondary antibody incubation in EveryBlot blocking buffer (BioRad, #12010020) (see Table S11 - Antibodies). The BioRad ChemiDoc imager was used to develop the membranes.

### CUT&RUN

CUT&RUN for histone marks, transcription factors and transcription co-factors was performed as previously described^60^. First, 10 µL of ConA beads (Polysciences, #86057) per reaction were activated by washing twice with 100 µL Bead Activation Buffer (20 mM HEPES (pH 7.9), 10 mM KCl, 1 mM CaCl_2_, 1 mM MnCl_2_) and finally resuspended in 10 µL Bead Activation Buffer.

Following, 250,000 and 1 million cells were used for the profiling of histone marks and transcription factors/co-factors, respectively. When applicable, cells were treated with 500 nM Entinostat – MS-275 (Selleckchem, #S1053) for 24 h prior to nuclear extraction. Cells were pelleted in FACS tubes at 400 × g for 3 min, washed with PBS and resuspended in 100 µL of NE Buffer (20 mM HEPES (pH 7.9), 10 mM KCl, 0.1% Triton X-100, 20% Glycerol, 1 mM MnCl_2_, 0.5 mM Spermidine) supplemented with 1× cOmplete EDTA-free protease inhibitor cocktail (Roche, #11836170001). Reactions were incubated on ice for 10 min and centrifuged at 400 × g for 3 min. Nuclear pellets were resuspended in 100 uL NE Buffer supplemented with 1× cOmplete EDTA-free protease inhibitor cocktail (Roche, #11836170001) and directly mixed with 10 µL activated ConA beads (Polysciences, #86057). Nuclei were allowed to bind ConA beads (Polysciences, #86057) for 10 min at RT. The supernatant was removed and the nuclei-bound beads were resuspended in 50 µL Antibody Buffer (20 mM HEPS (pH 7.5), 150 mM NaCl, 0.5 mM Spermidine, 0.02% digitonin, 2 mM EDTA, 1× cOmplete EDTA-free protease inhibitor cocktail (Roche, #11836170001)) containing 1 µL of the respective antibody (see Table S11 - Antibodies). Tubes were nutated at 4 °C overnight. The following day, beads were washed twice with 200 µL Digitonin Buffer (20 mM HEPS (pH 7.5), 150 mM NaCl, 0.5 mM Spermidine, 0.02% digitonin, 1× cOmplete EDTA-free protease inhibitor cocktail (Roche, #11836170001)) and resuspended in 50 µl of Digitonin Buffer with pAG-MNase (Institute of Molecular Pathology, VBC, Vienna, Austria). Beads were incubated for 30 min at RT and then washed twice with 200 µL Digitonin Buffer. Following, MNase was activated as beads were resuspended in 50 µL Digitonin Buffer supplemented with 2 mM CaCl_2_ and incubated for 30 min at 4 °C. The reaction was quenched upon addition of 33 µL of Stop Buffer (340 mM NaCl, 20 mM EDTA, 4 mM EGTA, 50 µg/mL RNaseA, 50 mg/mL Glycogen). Following incubation at 37 °C for 10 min, the supernatant was transferred to a fresh tube and DNA was purified using the NEB Monarch PCR &DNA Clean-up Kit (NEB, #T1030).

Libraries were prepared as previously descried (<u>Library Prep for CUT&RUN with NEBNext® Ultra™ II DNA</u> <u>Library Prep Kit for Illumina® (E7645) (protocols.io)</u>) using the NEBNext Ultra II DNA Library Prep Kit for Illumina (NEB, #E7645) and the NEBNext Multiplex Oligos for Illumina (Dual Index Set 1) (NEB, #7600). 2 ng of CUT&RUN DNA was used as input for library preparation using 1.5 µM adaptors, 14-16 PCR amplification cycles and a one-sided 0.9× size selection.

### CUT&RUN analysis

Sequencing reads (150 bp paired-end, NovaSeq 6000) were trimmed using TrimGalore (ersion 0.6.6)^61^ and aligned to human genome, hg38, using Bowtie2 (version 2.4.2)^62^ (--local, --very-sensitive, --no-mixed, --no-discordant, --dovetail). SAMtools (version 1.15.1)^63^ was used to fix mates of paired end reads, remove PCR duplicates of IgG sample, merge, sort and index bam files, while a CUT&RUN-specific blacklist^64^ was removed from the bam files using BEDtools (version 2.30.0)^65^.

DeepTools (version 3.5.1)^66^ served to build the genome coverage, to compute the matrix and plot the profile of histone marks, ARID5B, HDAC1 and HDAC2. Average signal intensities around ARID5B peaks (+/- 5 kb) were calculated directly from the matrix obtained from deepTools (version 3.5.1)^66^. Peaks of histone marks were called using SEACR (relaxed with IgG as control)^67^. For the calling of ARID5B, HDAC1 and HDAC2 peaks, IgG signal was first subtracted from the signal of the protein of interest using bdgcmp from MACS2 (version 2.2.5) (-m subtract)^68^. This was then used as an input file for callpeaks from MACS2 (version 2.2.5) (--cutoff 0.5, -l 200, -g 150; for ARID5B: --cutoff 0.4, -l 200, -g 150)^68^. For downstream analysis, BEDtools (version 2.30.0)^65^ was used to intersect peaks, SeqCode (version 1.0)^69^ and ChromHMM (version 1.24)^70^ were employed to determine the genomic localization of ARID5B peaks, HOMER (version 4.11.1)^71^ and ChIP Atlas^72^ were utilized to investigate TF motifs and binding on the regions of interest, and DiffBind (version 3.16.0)^73^ was run for differential binding analysis (DBA_EDGER, without IgG control, 0.8 ≥ log2FC, and FDR ≤ 0.05). Data was visualized using ggplot2 (version 3.5.1)^74^.

### Processing of publicly available ChIPseq data

Fastq files of H3K4me3 ChIPseq data in NALM6 (SRR708094) were trimmed using TrimGalore^61^ and mapped to the human genome, hg38, using Bowtie2 (version 2.4.2)^62^ (--local, --very-sensitive). SAMtools (version 1.15.1)^63^ was used to convert SAM to BAM files, sort BAM files, remove PCR duplicates and index BAM files. Finally, deepTools (version 3.5.1)^66^ were used to build the genome coverage, compute the matrix and plot the profile on ARID5B peaks. BAM files were also used as input in the ChromHMM (version 1.24)^70^ analysis, as discussed in the section above.

### RNA isolation and RT-qPCR

RNA was harvested from 2 million cells using the RNeasy Plus Mini Kit (Qiagen, #74134). 1 µg of RNA was used for cDNA synthesis using the iScript cDNA Synthesis Kit (BioRad, #1708891). qPCR was performed using the iQ5 SYBR Green Supermix (BioRad, #1708885) and ran in the CFX Connect Real Time System (BioRad). An initial denaturation at 95 °C for 3 min, followed by 40 cycles of 10 s denaturation at 95 °C and 30 s annealing and extension at 60 °C was used (see Table S11 - Oligonucleotides).

### mRNA-seq library preparation

RNA was isolated as described above and isent to Azenta (Leipzig, Germany) for QC and library preparation followed by next-generation sequencing (150 bp paired-end, NovaSeq 6000). When applicable, cells were treated with 500 nM Entinostat – MS-275 (Selleckchem, #S1053) for 24 h prior to RNA harvesting. Al mRNA-seq experiments were conducted using biological triplicates per condition.

### mRNA-seq analysis

Sequencing reads were trimmed using TrimGalore (version 0.6.6)^61^, aligned to the human transcriptome, hg38, using STAR (version 2.7.9)^75^ and counted with HTseq-count (version 0.11.2)^76,77^. DESeq2 (version 1.46.0)^78^ was used for differential gene expression (0.8 log2FC, basemean expression ≥30 and padj ≤ 0.05). Gene Set Enrichment Analysis (GSEA) (version 4.3.2)^79^ was run using the normalized read counts from DEseq2 (version 1.46.0) ^78^ of expressed genes (normalized read counts ≥ 30) with default settings and a gene set permutation type. pheatmap and ggplot2 (version 3.5.1)^74^ were used for data visualization.

### Statistical Analysis

R and the limma package^49^ were used to identify significantly enriched proteins in the interaction proteomics dataset. P-values were calculated using empirical Bayes moderation, followed by multiple testing correction using the Benjamini-Hochberg procedure. All other statistical analyses were conducted using GraphPad Prism (version 10.4.1). The Kolmogorov-Smirnov test was employed when comparing histone mark or co-factor signal intensities on ARID5B peaks, as the results are not expected to follow a normal distribution (non-parametric) and as the average occupancy on all of these sites (unpaired) was compared. Unpaired t-tests were performed when comparing gene expression between two conditions. One-way ANOVA followed by Tukey multiple comparison test was performed when comparing gene expression with more than two conditions. *P* ≤ 0.05 were considered statistically significant. \**P* ≤ 0.05, \*\**P* ≤ 0.01, \*\*\**P* ≤ 0.001, ns = not significant.

## Supplementary Figures

**Figure S1:**
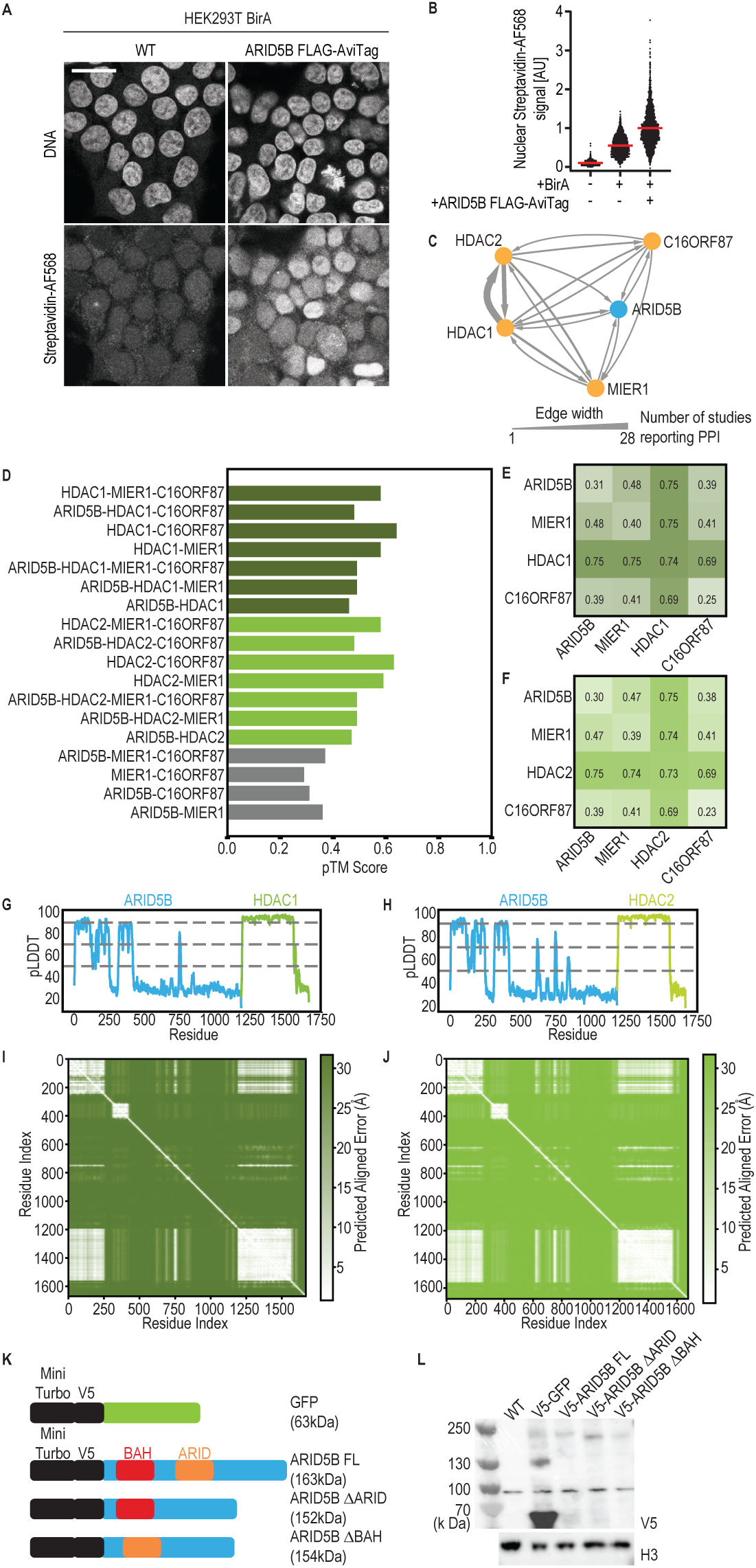
ARID5B interacts with transcriptional repressors. **a**. Representative immunofluorescence images of HEK293T cells stably expressing the E. Coli BirA biotin ligase, with or without expression of a murine ARID5B FLAG-AVI-tag cDNA construct. Cells were fixed for immunofluorescence and biotin deposition was assessed using Streptavidin conjugated to Alexa Fluor 568. DNA was stained with Hoechst 33342. Scale bar: 20 µm. **b.** Quantification of nuclear Streptavidin AF568 fluorescence by immunofluorescence for cells in **(a)** Wild type HEK293T cells were also used as a negative control to measure background staining. Dots represent the mean nuclear Streptavidin AF568 fluorescence of individual cells; red bars indicate the mean. Significance was tested using a two-tailed Mann–Whitney U-test. Number of cells analyzed (merge of n = 2 biological replicates): Wild type (n = 1255), BirA overexpression (n = 2032), BirA + ARID5B overexpression (n = 2672). **c.** Directional PPI-network of ARID5B co-repressor complex subunits obtained from publicly available interaction proteomics datasets. Edge arrows indicate the direction of interaction as reported in the database, while edge thickness corresponds to the number of associated publications. **d.** Predicted template modelling (pTM) scores from AlphaFold3 models for all predicted complex combinations. **e., f.** Chain-pair ipTM score matrices for AlphaFold3- predicted ARID5B-MIER1-C16ORF87 complexes with HDAC1 **(e)** or HDAC2 **(f)**. **g., h.** Per-residue predicted local distance difference test (pLDDT) scores for AlphaFold3-predicted ARID5B-HDAC1 **(g)** and ARID5B-HDAC2 **(h)** complexes. **i., j.** Predicted aligned error (PAE) heatmaps for ARID5B-HDAC1 **(i)** and ARID5B-HDAC2 **(j)** complexes. **k.** Schematic overview of the constructs used for co-immunoprecipitation experiments. GFP, full-length (FL) ARID5B and ARID or BAH domain deletion mutants (ΔARID or ΔBAH, respectively) containing a N-terminus MiniTurbo and V5 Tag were overexpressed in HAP1 cells expressing Cas9. **l.** Assessment of overexpression constructs for co-immunoprecipitation. Western Blot for V5 of protein lysates of HAP1 Cas9 and HAP1 Cas9 overexpressing V5-tagged GFP, full-length (FL) ARID5B and ARID or BAH domain deletion mutants (ΔARID or ΔBAH, respectively). Expected protein sizes are 63 kDa, 163 kDa, 152 kDa and 154 kDa for GFP, ARID5B FL, ΔARID and ΔBAH, respectively. H3 was used as a loading control.

**Figure S2:**
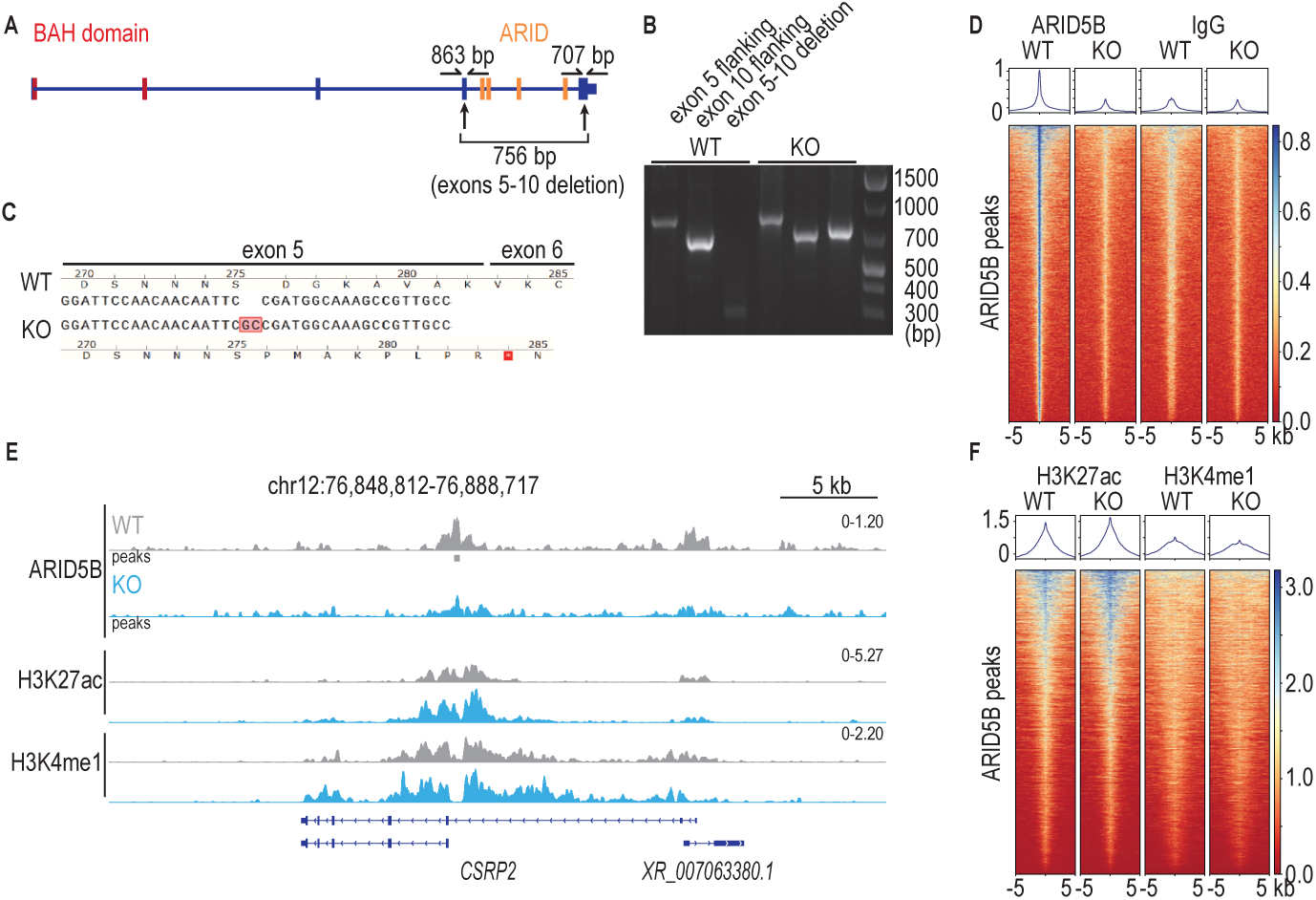
Genome-wide profiling of ARID5B. **a**. Scheme depicting the strategy for *ARID5B* KO. NALM6 WT cells were electroporated with RNPs with sgRNAs targeting *ARID5B* exons 5 and 10. Primers flanking the targeted regions were used to assess deletion of the exon 5-10 region as well as indels and point mutations. **b.** Genotyping of NALM6 WT and ARID5B KO cells. As expected, in NALM6 WT cells, PCR products are only observed for regions flanking the targeted *ARID5B* loci. ARID5B KO cells appear to be compound heterozygous, with one frameshift allele retaining exons 5 and 10 and the other allele presenting a deletion of the *ARID5B* exon 5-10 segment. **c.** Sanger sequencing of the targeted exon 5 in ARID5B KO cells reveals a 2 bp insertion, leading to a premature stop codon. **d.** Aggregate plots and heatmaps of ARID5B and IgG in NALM6 WT and ARID5B KO cells centered on the previously determined 4,200 ARID5B peaks. The coverage of two merged biological replicates per ARID5B and IgG per genotype was used for the analysis. **e.** Genome-wide coverage of ARID5B, H3K27ac and H3K4me1 in NALM6 WT and NALM6 ARID5B KO cells at the *CSRP2* locus. The coverage of two merged biological replicates per histone mark and ARID5B per genotype was used for the analysis. **f.** Aggregate plots and heatmaps of H3K27ac and H3K4me1 in NALM6 WT and ARID5B KO cells centered on the previously determined 4,200 ARID5B peaks. The coverage of two merged biological replicates per histone mark was used for the analysis.

**Figure S3:**
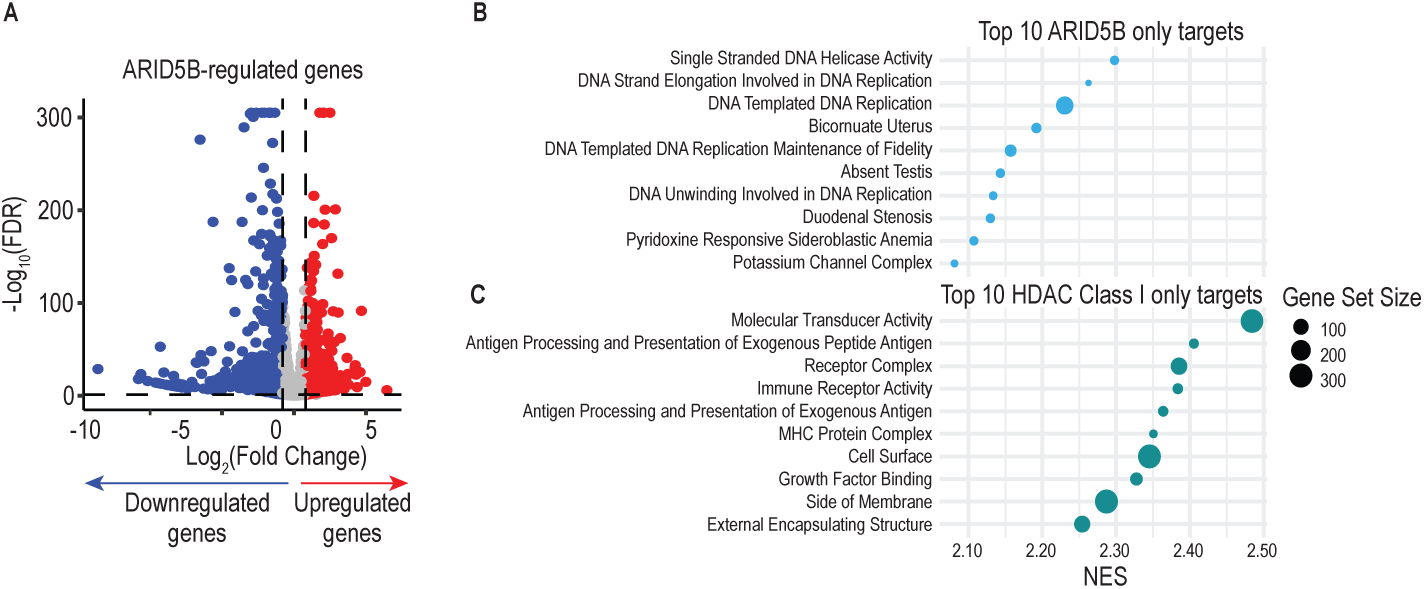
Pathways regulated by ARID5B and HDAC Class I. **a**. Volcano plot of genes regulated ARID5B KO in NALM6 cells. Highlighted in red and blue are up- and downregulated genes, respectively. Dotted lines indicate a log2FC threshold of larger or equal to 0.8 and a FDR threshold of smaller or equal to 0.05. The following thresholds were used for up- (log2FC ≥ 0.8, padj ≤ 0.05, cpm (KO) ≥ 30) and downregulated (log2FC ≤ -0.8, padj ≤ 0.05, cpm (WT) ≥ 30) genes. Biological triplicates per genotype and per treatment were used. **b.** Top 10 Gene Ontology terms enriched in gene set enrichment analysis (GSEA – C2 collection) of ARID5B KO NALM6 cells only. Depicted are the normalized enrichment scores (NES) and the number of genes contained in each gene ontology term. GSEA was run comparing NALM6 WT with ARID5B KO cells. Terms with an FDR smaller or equal to 0.25 were considered statistically significant. **c.** Top 10 Gene Ontology terms enriched in gene set enrichment analysis (GSEA - C2 collection) of Entinostat treated NALM6 cells only. Depicted are the NES and the number of genes contained in each gene ontology term. GSEA was run comparing vehicle-treated with Entinostat (MS-275)- treated cells. Terms with an FDR smaller or equal to 0.25 were considered statistically significant. NALM6 WT cells were treated with vehicle or 500 nM Entinostat (MS-275) for 24 h prior to mRNAseq experiment.

